# CuBlock: A cross-platform normalization method for gene-expression microarrays

**DOI:** 10.1101/2020.10.29.360198

**Authors:** Valentin Junet, Judith Farrés, José M. Mas, Xavier Daura

## Abstract

**Motivation:** Cross-(multi)platform normalization of gene-expression microarray data remains an unresolved issue. Despite the existence of several algorithms, they are either constrained by the need to normalize all samples of all platforms together, compromising scalability and reuse, by adherence to the platforms of a specific provider, or simply by poor performance. In addition, many of the methods presented in the literature have not been specifically tested against multi-platform data and/or other methods applicable in this context. Thus, we set out to develop a normalization algorithm appropriate for gene-expression studies based on multiple, potentially large microarray sets collected along multiple platforms and at different times, applicable in systematic studies aimed at extracting knowledge from the wealth of microarray data available in public repositories; for example, for the extraction of Real-World Data to complement data from Randomized Controlled Trials. Our main focus or criterion for performance was on the capacity of the algorithm to properly separate samples from different biological groups.

**Results:** We present CuBlock, an algorithm addressing this objective, together with a strategy to validate cross-platform normalization methods. To validate the algorithm and benchmark it against existing methods, we used two distinct data sets, one specifically generated for testing and standardization purposes and one from an actual experimental study. Using these data sets, we benchmarked CuBlock against ComBat (Johnson *et al*., 2007), YuGene (Lê Cao *et al*., 2014), DBNorm (Meng *et al*., 2017), Shambhala (Borisov *et al*., 2019) and a simple log_2_ transform as reference. We note that many other popular normalization methods are not applicable in this context. CuBlock was the only algorithm in this group that could always and clearly differentiate the underlying biological groups after mixing the data, from up to six different platforms in this study.

**Availability:** CuBlock can be downloaded from https://www.mathworks.com/matlabcentral/fileexchange/77882-cublock

**Supplementary information:** Supplementary data are available at *bioRxiv* online.

## 1 Introduction

Since the first whole-genome microarray study of gene expression was published in 1997 (Schena *et al*., 1995; Lashkari *et al*., 1997), high-throughput gene-expression microarrays have been a standard in many experimental designs in biological and biomedical research. Although their use is being replaced by next-generation sequencing techniques such as RNA-Seq (Nagalakshmi *et al*., 2008), the large amounts of microarray data relevant to an equally large variety of biological and biomedical problems and available in public databases constitutes a valuable resource that will remain in use for many years. The potentiality of resources such as the Gene Expression Omnibus (GEO) as sources of Real-World Data (RWD) —data derived from a number of sources, outside the context of Randomized Controlled Trials (RCTs), and associated with outcomes in an heterogeneous patient population (Berger *et al*., 2017)–may in fact boost the use of the wealth of available microarray data in the near future. The importance of RWD as a complementary information source in drug-evaluation studies is based on the observation that data from RCTs does not always match results from observational studies (Trotta, 2012), mostly owing to the limited number of RCT patients, their over-monitoring and the limited follow-up time. Thus, certain adverse drug reactions or lack-of-efficacy problems are hidden until RWD studies are performed, and drug administrations have come to encourage the extraction of information from sources complementary to RCTs to increase evidence around treatments (Sherman *et al*., 2017; Food and Drug Administration, 2018). A problem of RWD is that it tends to be highly heterogeneous, thus requiring careful analysis and statistical treatment (Berger *et al*., 2017; Bartlett *et al*., 2019). Translated to the context of this study, microarray data relevant to a particular problem will often originate from different laboratories and experiments, possibly using different microarray platforms (Bumgarner, 2013) and almost certainly obtained in different batches. In order to make a sensible use of such an heterogeneously sourced data, a data-normalization step is required before data analysis. Normalization can be relatively straightforward when dealing with different batches of a same experiment using the same biological samples, platform and operator, but gets increasingly complex as different operators, platforms and sample sources are introduced. This often leads to studies discarding part of the available data, which could otherwise be used to increase the chances of discovery of meaningful patterns or improve their statistics.

A main source for sample differences arising from systematic biases is the mixing of data from different microarray platforms. Unfortunately, most standard and widely used normalization methods are applicable to or have been developed for the single-microarray-platform context (Rudy and Valafar, 2011), making them generally inappropriate for the cross-study analysis of existing data sets. On the other hand, most existing cross-platform normalization methods, such as ComBat (Johnson *et al*., 2007), XPN (Shabalin *et al*., 2008) or DWD (Benito *et al*., 2004), require the data from different platforms to be normalized together —XPN and DWD were in fact developed for pairwise cross-platform normalization. For large data sets, normalizing platforms together can be restrictive. In addition to involving the normalization of a large joined data set, the eventual addition of new microarray data requires global renormalization. This led to the more recent development of sample-wise, cross-platform normalization methods such as SCAN (Piccolo *et al*., 2012), YuGene (Lê Cao *et al*., 2014), DBNorm (Meng *et al*., 2017) —which can operate sample or platform wise– and Shambhala (Borisov *et al*., 2019). SCAN performs a sample-wise normalization assuming a double Gaussian mixture distribution. It was, however, specifically designed for Affymetrix platforms, thereby restricting its general use. The other distribution-based normalization method, DBNorm, scales the data distributions from the individual microarrays to a common form, which does not need to be predetermined (e.g. the distribution from a reference microarray). As a downside, it is very slow. On the other end, YuGene uses a simple transform that assigns a modified cumulative proportion value to each measurement, making the normalization very fast. Finally, Shambhala uses a harmonization method that transforms each profile so that it approaches the output of a chosen golden-standard platform.

Although the number of normalization methods proposed in the literature is large, to our knowledge there are no other major cross-platform normalization methods that can be applied to gene-expression microarrays in a platform agnostic way and that have been tested and validated as such. To enable systematic studies involving the download of microarray data from databases (possibly at different times) and its normalization and storage for later retrieval, allowing anon-linear use of the data—for example, in successive analysis incorporating different amounts of data as available or necessary, it is essential that a downloaded microarray set need not be normalized more than once. Here, we introduce a novel cross-platform normalization method fulfilling all these conditions. The algorithm is called CuBlock, which stands for Cubic approximation by Block. We validate its performance using various metrics and compare it to five methods that can be used in a cross-platform context, namely, the log_2_ transform of raw data, ComBat, YuGene, DBNorm and Shambhala. Although ComBat does not normalize platforms separately —it was not devised as a normalization method but as one to adjust the data for batch effects, we introduced it in this benchmark set as a popular first choice for cross-platform normalization and a frequent benchmark standard for other methods (Walsh *et al*., 2015; Irigoyen *et al*., 2018). Overall, CuBlock shows the best performance in this group.

## 2 Methods

In this section we introduce the data sets used for the validation of CuBlock and describe the data preprocessing approach and the methods used for benchmarking and validation.

### 2.1 The data sets

We selected two benchmark data sets previously used in similar studies (Rudy and Valafar, 2011; Borisov *et al*., 2019). The first set (here called the reference data set) originates from projects MAQC (MAQC-I) (MAQC Consortium, 2006) and SEQC/MAQC-III (SEQC/MAQC-III Consortium, 2014), which made use of reference RNA samples to assess repeatability of gene-expression microarray data within a specific site, reproducibility across multiple sites and comparability across multiple platforms. The second set (here called the experimental data set) originates from a study trying to assess profile differences of human spermatozoal transcripts from fertile and teratozoospermic males (Platts *et al*., 2007). The use of these two data sets allows us to assess, independently, effects from technical replicates (same biosample, analyzed in different labs with repetition) and biological replicates (different biosamples corresponding to a same condition).

#### 2.1.1 The reference data set

The data of this set are accessible in GEO with accession numbers GSE5350 (MAQC-I) and GSE56457 (MAQC-III), respectively. The set contains microarray gene-expression data corresponding to four titration pools from two distinct reference RNA samples: (*A*) Stratagene’s Universal Human Reference RNA pool; (*B*) Ambion’s Human Brain Reference RNA pool; (*C*) pool with an *A:B* ratio of 3:1; (*D*) pool with an *A:B* ratio of 1:3. These biosamples had been analyzed using different platforms and in different sites, as described (MAQC Consortium, 2006; SEQC/MAQC-III Consortium, 2014). Following the work from Rudy and Valafar(2011) and Borisov et al. (2019), we selected data from six of the platforms (between parentheses, data-set identifier in this study, GEO platform ID and project of origin):

- Affymetrix Human Genome U133 Plus 2.0 Array (AFX, GPL570, MAQC-I): 3 experiments (sites) (AFX_1 to AFX_3), with 4 biosamples (*A-D*) per experiment and 5 replicates per biosample (60 samples)
- Agilent-012391 Whole Human Genome Oligo Microarray G4112A (AG1, GPL1708, MAQC-I): 3 experiments (AG1_1 to AG1_3), with 4 biosamples (*A-D*) per experiment and 5 replicates per biosample (60 samples)
- Illumina Sentrix Human-6 Expression BeadChip (ILM, GPL2507, MAQC-I): 3 experiments (ILM_1 to ILM_3), with 4 biosamples (A-D) per experiment and 5 replicates per biosample (59 valid samples)
- Illumina Human HT-12V4.0 Expression Beadchip(HT12, GPL10558, MAQC-III): 2 experiments (ILM_COH and ILM_UTS), with 4 biosamples (*A-D*) per experiment and 3 replicates per biosample (24 samples)
- GeneChip^®^ PrimeView™ Human Gene Expression Array (PRV, GPL16043, MAQC-III): 1 experiment (AFX_USF_PRV), with 4 biosamples (*A-D*) and 4 replicates per biosample (16 samples)
- Affymetrix Human Gene 2.0 ST Array (HUG, GPL17930, MAQC-III): 1 experiment (AFX_USF_HUG), with 4 biosamples (*A-D*) and 4 replicates per biosample (16 samples)

Note that in the MAQC-I study the following microarrays from AG1 were discarded as outliers after the Agilent’s Feature Extraction QC Report: AG1_1_A1, AG1_2_A3, AG1_2_D2, AG1_3_B3. Since the data for these microarrays is nevertheless deposited and we wanted our analysis to be as independent as possible of platform-dependent data-preprocessing steps, we considered also their inclusion. To this end, we evaluated the correlation of the data between all AG1 samples and observed that the “outliers” are highly correlated to the non-outliers of the same experiment and of the other two experiments (about 0.97 in both cases). A dimension reduction of the raw data showed also no outliers. We therefore decided to include these four microarrays in the data set.

#### 2.1.2 The experimental data set

This data set contains spermatozoal RNA samples from normally fertile (*N*) and heterogeneously teratozoospermic (*T*) subjects and is accessible in GEO with accession number GSE6969. The samples had been analyzed on three different platforms (between parentheses, data-set identifier in this study and GEO platform ID):

- Affymetrix Human Genome U133 Plus 2.0 Array (AFF, GPL570): 13 independent biosamples of type *N* and 8 of type *T*
- Illumina Sentrix Human-6 Expression BeadChip (ILL1, GPL2507): 5 independent biosamples of type *N* and 8 of type *T*
- Illumina Sentrix HumanRef-8 Expression BeadChip (ILL2, GPL2700): 4 independent biosamples of type *N* and 6 of type *T*

### 2.2 Data processing

To make the analysis as platform agnostic as possible, we took the image-processed raw intensities for all non-control probes and disregarded any platform-dependent background-signal correction such as that provided by mismatch probes in Affymetrix platforms. CEL files for Affymetrix and txt files for the other platforms were used. Probes with invalid intensities (NaN) in data sets HT12 and HUG were ignored. For Affymetrix microarrays, the intensities of probes constituting a probe set were averaged. CuBlock normalization was then applied to the log_2_ transform of the probe intensities, for each platform separately. Since probes vary among the different microarray platforms, the normalized data sets were then transformed from the probe level to the protein level by mapping probes to UniProtKB accession numbers (ACs) and keeping only those probes that map to an AC present in all platforms. Each selected AC was then assigned an intensity equal to the average of the normalized intensities of associated probes in the given microarray.

To benchmark CuBlock against established normalization methods applicable in a generic cross-platform context, we compared it to a simple log_2_ transform and to the methods ComBat (Johnson *et al*., 2007), YuGene (Lê Cao *et al*., 2014), DBNorm (Meng *et al*., 2017) and Shambhala (Borisov *et al*., 2019). YuGene and DBNorm were applied following the same procedure used for CuBlock, i.e., normalization of the log_2_ transform of the probe intensities and successive mapping to ACs. ComBat requires all microarrays to be normalized together, which implies their merging before normalization. Therefore, in this case the mapping to UniProtKB ACs and selection of ACs present in the different platforms was performed after log_2_ transform and before ComBat normalization. We note that DBNorm allows normalization per sample and per platform. We performed both, but show only the results obtained with samplewise normalization since they are better. Comparison to Shambhala was done only for the data sets AFX, AG1 and ILM from the reference data set, since Shambhala-normalized data for these sets has been already reported by the authors as supplementary data to Borisov et al. (2019). DBNorm was only used on the experimental data set, as the calculations turned out to be forbiddingly slow. To perform the calculations we used the R package sva for ComBat(https://bioconductor.org/packages/release/bioc/html/sva.html) and the packages provided by YuGene (https://cran.r-project.org/web/packages/YuGene/index.html) and DBNorm (https://github.com/mengqinxue/dbnorm) authors in the respective papers. Calculations with these programs were performed with default settings. For DBNorm, in order to reproduce the general case (e.g. this study), in which a reference microarray cannot be straightforwardly selected, we used the option of normalization into a normal distribution.

### 2.3 Comparison and validation methods

To validate and compare the cross-platform normalization methods evaluated in this study we used the methodology described below. The objective was to increase the sensitivity, i.e. the identification of true biological differences, while minimizing platform and various kinds of replica effects. All validation methods were applied on a subset of 500 proteins that best distinguish two given biological groups.

To select the 500 proteins we first performed a differential analysis on the normalized data, for all platforms in the reference or experimental data set. To this end, we performed Welch’s t-test to evaluate, for each protein, the difference between the associated mean intensities in units of uncertainty (the t-statistic) in two biological groups, *A* and *B* from the reference data set (total of 16624 proteins) or *N* and *T* from the experimental data set (total of 16937 proteins). Note that we deliberately avoid considerations on whether the data sets meet the requirements of the *t*-test, since we used the test simply to identify the 500 proteins with largest separation of group means per uncertainty unit, that is, with lowest associated *p*-values, irrespective of the error in the *p*-value and, therefore, of its valid interpretation as a probability. Although for such purpose we do not require the calculation of FDR-adjusted *p*-values (*q*-values) (Storey, 2002), since they conserve *p*-value ranking, we did obtain them and show corresponding ROC-like curves (the cumulative distribution function of the *q*-values) in Figure S1 (Supplementary information). We decided to select a fixed number of proteins, rather than proteins with a *p*- or *q*-value below a given arbitrary threshold, to enable the comparison of methods using data sets of equal and reasonably large dimensionality. We note, nevertheless, that the 500-protein cut corresponds to an FDR well below 10^−2^ (Figure S1). The differential analysis was performed with the MATLAB function *mattest*.

#### 2.3.1 Silhouette plot

Silhouette plots are graphical displays of data partitions (Rousseeuw, 1987), where clusters are represented by so-called *silhouettes* generated by comparison of cluster tightness and separation. The method assigns a silhouette value between −1 and 1 to each element of a cluster, indicating if the element is well clustered (value close to 1), lies between two or more clusters (close to 0) or is likely misclassified (close to −1). The silhouette plot is then generated by representing the values for all elements as bars, for the different cluster partitions. We computed three silhouette plots for the reference data set: one identifying clusters with platforms, one where the data was assigned to groups *A*∪*C* and *B*∪*D* and one where the partitioning was represented by sets *A, B, C* and *D*. For the experimental data set, silhouette plots based on platform partitioning and *T vs. N* partitioning were computed. The MATLAB function *silhouette* was used to compute the silhouette plots.

#### 2.3.2 *t*-SNE dimension reduction

*t*-SNE (Maaten and Hinton, 2008) is a stochastic dimension-reduction method aimed to preserve the local structure of data (keeping the lowdimensional representation of very similar data points close together) while retaining essential traits of global structure. It analyzes the neighborhood of the data points by calculating pairwise conditional probabilities representing their similarity. The method then tries to find a low-dimensional representation that minimizes the difference between the high-dimensional and low-dimensional conditional probabilities. The parameter controlling the number of neighbors is called perplexity, and is typically given values between 5 and 50. Due to its stochastic nature and the dependence on the chosen perplexity parameter, the algorithm may converge to irrelevant solutions. We thus performed 10 runs for each of a number of perplexity values and selected the one producing the most consistent biological partitioning according to the average silhouette values. For the reference data set we used perplexity values from 5 to 50, in increments of 5, and selected the representation giving the best clustering relative to sets *A*, *B*, *C* and *D*. For the experimental data set, we used perplexity values 5,10 and 15 and selected the representation giving the best clustering relative to sets *T* and *N*. The MATLAB function *tsne* was used to perform the *t*-SNE dimension reduction.

#### 2.3.3 Dendrogram

We performed a hierarchical clustering analysis using the Euclidean distance as metric and the arithmetic mean as linkage criterion, and represented the resulting cluster hierarchy as a dendrogram. To assess the significance of the clusters, we applied multiscale bootstrap resampling as provided in the R package *pvclust* (Suzuki and Shimodaira, 2006). By default, this package considers 10 relative bootstrap sample sizes (bootstrap sample size divided by total sample size), from 0.5 to 1.4, with 1000 resamplings per sample size, leading to a total of 10000 bootstrap resamples. The package provides two statistics to estimate the significance of the obtained clusters: the bootstrap probability (BP) or frequency (expressed as percentage) of observation of a given cluster in the bootstrap resamples, and the approximately unbiased *p*-value (AU), an unbiased version of BP. More details on multiscale bootstrap resampling can be found in Shimodaira (2004). We plot the dendrograms using the R package *dendextend*.

#### 2.3.4 SVM classification

The goal of this analysis was to assert whether relevant patterns can be found using the data from only one platform. Support vector machines (SVM) (Cortes and Vapnik, 1995) are binary classifiers applicable to problems that are reducible to a binary outcome, such as the *T* and *N* phenotypes in our experimental data set. We trained a linear support vector machine model for each platform using the following approach. To reduce feature-vector dimensionality, where dimensions are proteins (more specifically their microarray-derived intensities), while retaining the capacity to asses how well the 500 proteins separate the *T* and *N* populations, the training was performed six times, starting with dimension 5 and increasing it up to dimension 10. For each of 1000 runs with a given dimensionality, we selected randomly from the 500 protein set as many proteins as dimensions, extracted the corresponding data from sets *T* and *N*, trained a linear SVM model for each platform, separately, and tested it on the other two platforms. This led to a total of 6000 models per platform. For each platform, we calculated the mean and standard deviation of different classification scores over the 6000 models, namely, Accuracy, Matthews Correlation Coefficient (MCC) (Boughorbel *et al*., 2017), Balanced Accuracy and Area Under the ROC Curve (AUC) (Fawcett, 2006). The MATLAB function *svm* was used to train the SVM models.

## 3 Algorithm

The main idea of CuBlock is to partition probes into clusters and, for each sample and probe cluster (i.e. for each data block), transform the data by a procedure that involves the fitting of a cubic polynomial to a mapped distribution of points. A pseudo code of the CuBlock algorithm is described in Figure 1. It calls two additional algorithms with pseudo codes provided in Figures S2 and S3 (Supplementary information).

**Fig. 1.**
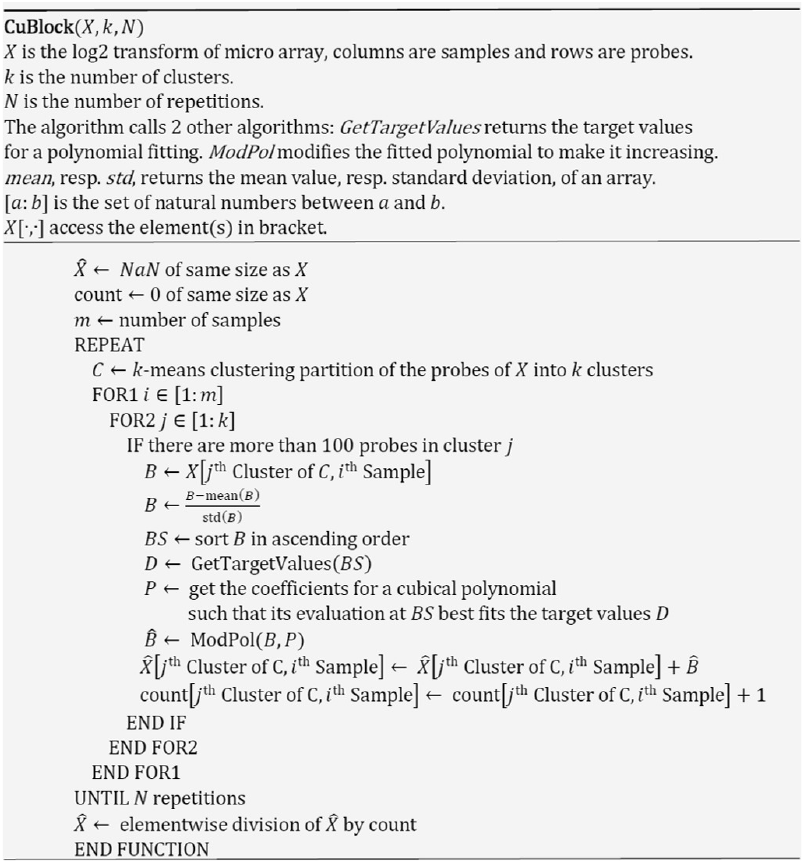
Pseudocode describing the CuBlock algorithm (see description in Section 3).

The input matrix *X* contains the log_2_ transform of the gene-expression microarray intensities, where columns are samples and rows are probes. CuBlock makes use of the *k*-means clustering algorithm (Lloyd, 1982) to partition probes in the space defined by the samples —a probe data point is a vector of probe intensities of dimension equal to the number of samples– and is applied per platform, i.e. the *k*-means clustering is performed for all samples of a given platform. *k*-means is an iterative algorithm that tries to partition the data into a predefined number *k* of non-overlapping clusters, starting from a random initialization of their centroids. Because of the random initialization, clusters from different runs may differ, and the core part of the CuBlock algorithm is repeated several times for different solutions of *k*-means. Thus, input parameters *k* and *N* in Figure 1 refer to the chosen number of *k*-means clusters and repetitions, which in this work took values of 5 and 30, respectively. The CuBlock algorithm finds first a probe-cluster partition in the space defined by the samples and then applies its normalization scheme to data blocks defined as those (log_2_) probe-intensity values from a sample that belong to a given cluster. Therefore, for *k* clusters and *m* samples we have a total of *k* · *m* blocks. The advantage of the normalization by block is that it decomposes the distribution of probe intensities of a sample into its different block distributions, according to similarities between probes found by the clustering algorithm in the space of all samples. These different distributions will enable the emergence of different patterns present in the data. Instead, if the blocks were selected at random or the whole sample was used, the normalization method would estimate parameters based on a unique distribution, masking these different patterns. Although we initially determine the probe clusters using all samples, we then normalize sample by sample to reduce the dependence of the normalization on the full sample collection. The strategy of normalization by block is similar to that used by the cross-platform normalization method XPN (Shabalin *et al*., 2008). However, XPN defines probe clusters and sample clusters with two independent application of *k*-means (one on the input matrix and the other on its transpose) and blocks are then constituted by all possible combinations of one sample cluster with one probe cluster.

For each block, and each of the N repetitions of the *k*-means clustering, CuBlock fits a cubic polynomial to a mapped set of points symmetrically distributed between −1 and 1 with density increasing toward zero. This is performed in four steps, as shown in Figure 1. First, the block data is linearly transformed to z-scores (zero mean and unit standard deviation). These are then used as input values of a mapping function whose output values will be used to fit the cubic polynomial, as described in the pseudocode shown in Figure S2. The mapping associates the sorted values present in the block to an equal number of equidistant points between −1 and 1, and takes these new points to an uneven power in order to have their distance decrease as they approach zero from either side (Figure S4). The exact uneven power will determine how slow is the growth of the points around zero, and is selected such that, on average, the values of the block that are within standard deviation, i.e. the block values between −1 and 1, are mapped to a value smaller than 0.1 (Figure S5). The algorithm tries uneven powers between 3 and 21 and the first one that fulfills the criterion is selected. Next, the algorithm finds the coefficients of a cubic polynomial that, when evaluated on the sorted block data (input values), best fits the output values from the mapping function (Figure S4). We chose to fit a cubic polynomial instead of a higher degree one to avoid overfitting. Polynomial coefficients were obtained with the MATLAB function *polyfit*, with degree 3.

If the block data is not symmetric or contains many outliers, a cubic polynomial will produce a poor fit. Thus, the polynomial will increase along the symmetric part of the block and decrease as it reaches the outliers (Figure S6). Despite leading to a poor fit, this feature can be used to identify asymmetry issues and outliers. When this is the case and decreasing values are identified after evaluating the polynomial on the block data, the decreasing values are corrected in order to preserve data sorting upon normalization. Roughly, the correction equates the decreasing values to the last increasing value (after an increasing section) or to the last decreasing value (before an increasing section). The precise corrections are described in Figure S6, and the cases where the cubic polynomial might decrease are considered in Figure S3.

Figure 2 shows the histograms of different samples before and after normalization. While before normalization the samples follow clearly different distributions, after normalization the distributions are much more homogeneous. We note that before normalization the distributions are clearly platform dependent (compare A and C, which correspond to the same biosample but different platforms, and B and C, which correspond to different biosamples and the same platform). This effect is remarkably corrected after normalization.

**Fig. 2.**
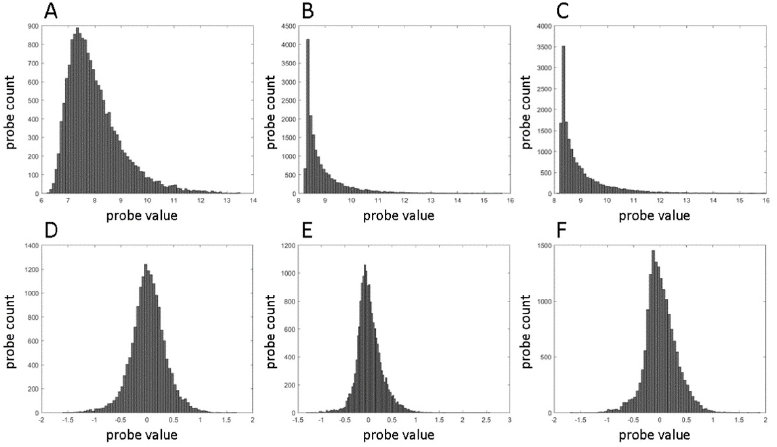
Example histograms for samples from different platforms. A-C: histograms before CuBlock normalization and after log_2_ transform. D-F: histograms after CuBlock normalization. A, D: biosample *A*, platform AFX; B, E: biosample *B*, platform AG1; C, F: biosample *A*, platform AG1.

## 4 Results and Discussion

The algorithm described in the previous section was applied to the data introduced in Section 2.1 after preprocessing (see Section 2.2), and the results were compared to those obtained with other normalization methods as explained in Section 2.3. In this section we will present and discuss these results.

### 4.1 Reference data set: six platforms

Figures 3 and 4 and Figure S7 show the results obtained with the different normalization methods using the dendrogram, silhouette and *t*-SNE analyzes, respectively. The three validation methods show that CuBlock and ComBat separate very clearly the biological groups *A*, *B*, *C* and *D* (except for a couple of *A* points in ComBat’s case). ComBat tends to produce tighter but less cleanly separated clusters for these four groups, as illustrated by both the *t*-SNE (Figure S7C) and silhouette (Figure 4C) plots. CuBlock is the only method that clusters the biological groups *A* and *C*, and *B* and *D* together in the dendrogram plot (Figure 3A), and this is also underlined in the corresponding silhouette plot in Figure S8A, showing high and homogeneous silhouette values. On the contrary, log_2_, ComBat and YuGene tend to cluster *C* with *D* (Figures 3B-D). In fact, log_2_ and YuGene have difficulties to separate this two groups at all. Figures S8 and S9 show silhouette plots using the groups *A*∪*C* and *B*∪*D* and the platforms as given clusters, respectively. We note that even though CuBlock puts emphasis on the biological differences and Figure S9A indicates weak platform clusters, both the *t*-SNE (Figure S7A) and dendrogram (Figure 3A) plots show that, within each of the *A, B, C, D* clusters, the samples are subclustered by platform. As can be seen in these Figures, ComBat mixes the data from the different platforms best, while YuGene and log_2_ are, in this order, worst at mixing platform data.

**Fig. 3.**
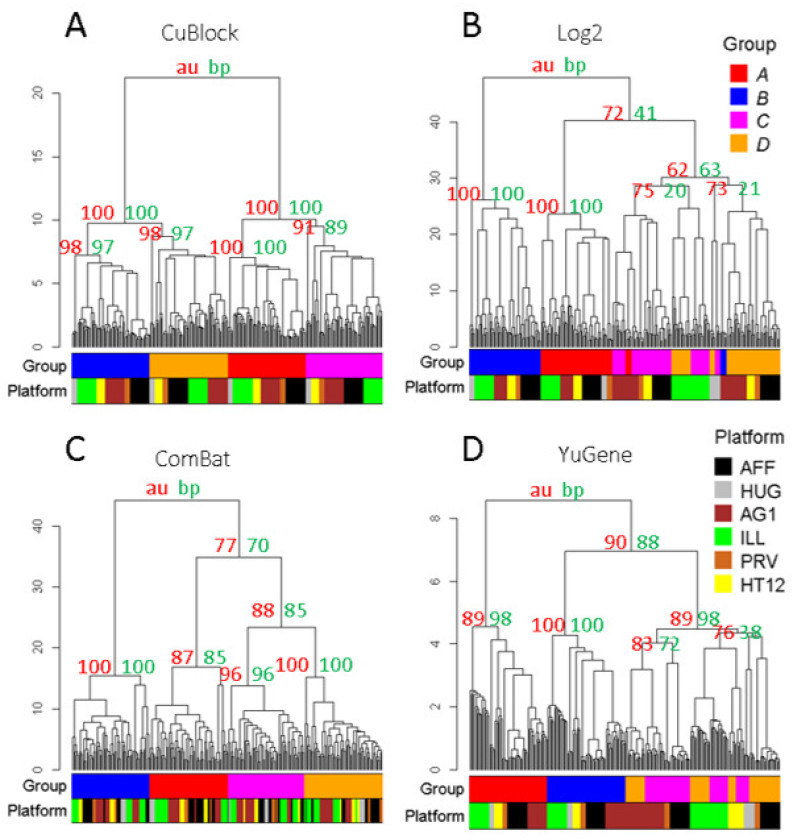
Dendrogram analysis of the reference data set (six platforms) after normalization with CuBlock (A), log_2_ (B), ComBat (C) and YuGene (D). Color bars under the dendrograms indicate the biological group and platform corresponding to each leaf; the BP (green) and AU (red) values (see Section 2.3.3) for some selected clusters are indicated at the origin of the branches.

**Fig. 4.**
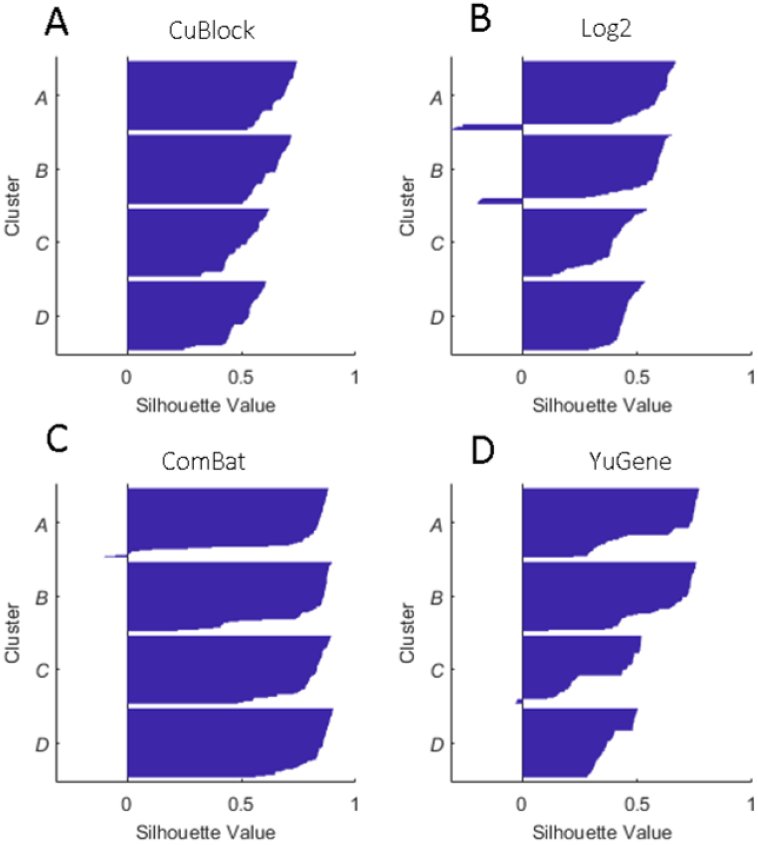
Silhouette plots of the reference data set (six platforms) after normalization with CuBlock (A), log_2_ (B), ComBat (C) and YuGene (D), using the groups *A, B, C* and *D* as given clusters. Mean silhouette index (SI) values: A: 0.66 (*A*), 0.62 (*B*), 0.51 (*C*) and 0.50 (*D*); B: 0.51 (*A*), 0.49 (*B*), 0.38 (*C*) and 0.44 (*D*); C: 0.72 (*A*), 0.78 (*B*), 0.79 (*C*) and 0.83 (*D*); D: 0.61 (*A*), 0.63 (*B*), 0.35 (*C*) and 0.39 (*D*).

### 4.2 Reference data set: three platforms

To compare the results from CuBlock and Shambhala (the latter reported by Borisov *et al*. (2019) for the same data set), we also performed the analysis for the three-platform subset used by the authors of Shambhala, namely AFX, ILM and AG1. They had concluded that Shambhala separates well *A*∪*C* from *B*∪*D* but not *A* from *C* or *B* from *D*. Using our selection of 500 proteins that best distinguish *A* from *B*, when looking at the results for Shambhala in Figure S10B,D we observe that, while *A*∪*C* forms a relatively clear cluster, all the *B*∪*D* points from AG1 samples are clustered with *A*∪*C*, making *B*∪*D* a well defined cluster only for AFX and ILM. As illustrated by the *t*-SNE and dendrogram plots and by the negative silhouette values in Figure S10F, Shambhala does also not distinguish *A, B, C* and *D* from each other well. The results for CuBlock in Figure S10A,C,E show the same features already discussed in Section 4.1 using the data for six platforms.

### 4.3 Experimental data set

Figures 5 and 6 and Figure S11 show the results obtained for the human sperm data set, after normalization with CuBlock, log_2_, ComBat, YuGene and DBNorm. CuBlock is the only normalization method that significantly distinguishes the two biological groups, *T* and *N*. The dendrogram plots in Figures 5 and *t*-SNE plots in Figure S11 show that the other methods tend to misclassify some samples, in particular the ones from platform ILL2. Similarly to the results for the reference data set (Section 4.1), CuBlock tends to sort the samples by platform within the clusters *T* and *N* (except for one *T* sample from ILL2). To investigate whether patterns that are found using one platform can be extrapolated to the other platforms, we performed a SVM classification test as described in Section 2.3.4. The results are shown in Table 1. In all cases, CuBlock outperforms the other methods. It is also worth noting that no matter which platform is used for the training based on CuBlock, the results are always very similar. To a lesser extent, this is also true for DBNorm. However, using log_2_, ComBat and YuGene, training with ILL2 gives worst results than with the other platforms, probably due to the fact that this platform constitutes a better defined cluster, as shown in the silhouette plots in Figure S12.

**Fig. 5.**
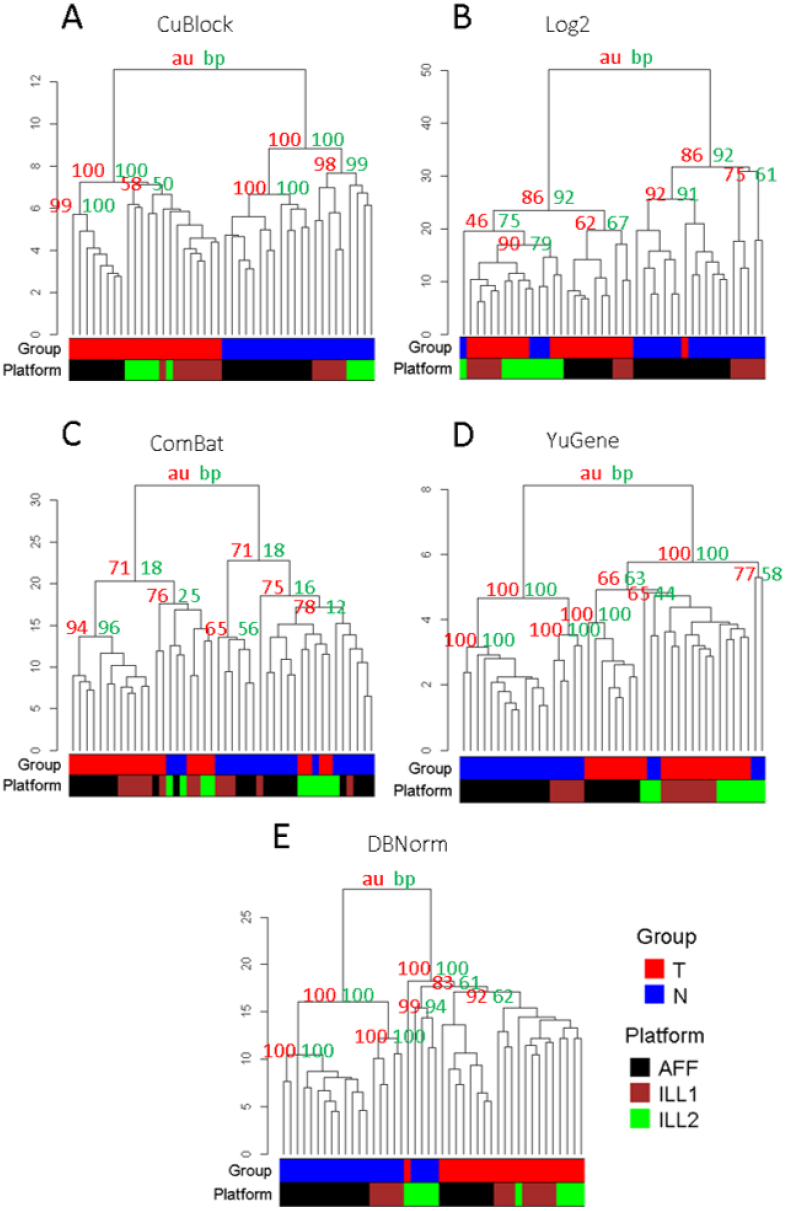
Dendrogram analysis of the experimental data set after normalization with CuBlock (A), log_2_ (B), ComBat (C), YuGene (D) and DBNorm (E). Color bars under the dendrograms indicate the biological group and platform corresponding to each leaf; the BP (green) and AU (red)values (see Section 2.3.3) for some selected clusters are indicated at the origin of the branches.

**Fig. 6.**
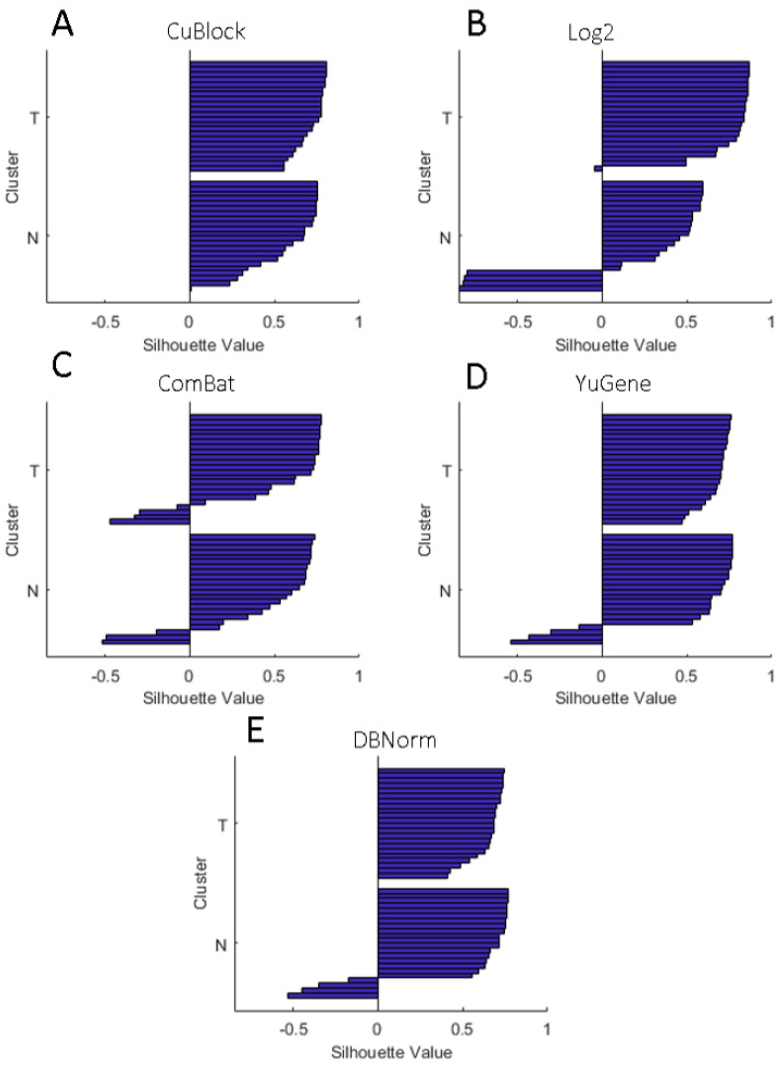
Silhouette plots of the experimental data set after normalization with CuBlock (A), log_2_ (B), ComBat (C), YuGene (D) and DBNorm (E) using the groups *T* and *N* as given clusters. Mean silhouette index (SI) values: A: 0.72 (*T*), 0.57 (*N*); B: 0.75 (*T*), 0.23 (*N*); C: 0.48 (*T*), 0.45 (*N*); D: 0.67 (*T*), 0.51 (*N*); E: 0.65 (*T*), 0.51 (*N*).

**Table 1.** SVM classification scores (see Section 2.3.4)

| Training platform | Accuracy | MCC | Balanced Accuracy | AUC |
| --- | --- | --- | --- | --- |
| CuBlock |  |  |  |  |
| AFF | 0.88 ± 0.08 | 0.76 ± 0.17 | 0.86 ± 0.10 | 0.98 ± 0.03 |
| ILL1 | 0.92 ± 0.06 | 0.85 ± 0.12 | 0.91 ± 0.07 | 0.99 ± 0.02 |
| ILL2 | 0.86 ± 0.16 | 0.74 ± 0.32 | 0.86 ± 0.17 | 0.99 ± 0.04 |
| log<sub>2</sub> |  |  |  |  |
| AFF | 0.77 ± 0.07 | 0.52 ± 0.16 | 0.71 ± 0.08 | 0.70 ± 0.11 |
| ILL1 | 0.80 ± 0.06 | 0.64 ± 0.11 | 0.81 ± 0.06 | 0.81 ± 0.04 |
| ILL2 | 0.44 ± 0.16 | -0.13 ± 0.35 | 0.46 ± 0.16 | 0.24 ± 0.31 |
| ComBat |  |  |  |  |
| AFF | 0.73 ± 0.11 | 0.53 ± 0.20 | 0.76 ± 0.10 | 0.90 ± 0.08 |
| ILL1 | 0.76 ± 0.08 | 0.57 ± 0.15 | 0.77 ± 0.08 | 0.90 ± 0.06 |
| ILL2 | 0.61 ± 0.30 | 0.22 ± 0.61 | 0.61 ± 0.30 | 0.63 ± 0.36 |
| YuGene |  |  |  |  |
| AFF | 0.81 ± 0.07 | 0.63 ± 0.15 | 0.77 ± 0.08 | 0.91 ± 0.06 |
| ILL1 | 0.87 ± 0.08 | 0.74 ± 0.16 | 0.87 ± 0.09 | 0.96 ± 0.04 |
| ILL2 | 0.55 ± 0.07 | 0.07 ± 0.17 | 0.53 ± 0.07 | 0.97 ± 0.11 |
| DBNORM |  |  |  |  |
| AFF | 0.81 ± 0.07 | 0.62 ± 0.16 | 0.77 ± 0.09 | 0.91 ± 0.07 |
| ILL1 | 0.86 ± 0.06 | 0.76 ± 0.09 | 0.87 ± 0.05 | 0.95 ± 0.04 |
| ILL2 | 0.79 ± 0.13 | 0.61 ± 0.24 | 0.77 ± 0.13 | 0.96 ± 0.11 |

Mean and standard deviation over the 6000 models per platform and method.

### 4.4 Reference data set: missing biological groups

CuBlock is a method that works with the actual distribution of the data without making any assumption on its shape. It is, in that sense, dependent on the actual differences found in the input data. To test this dependence, we analyzed again the reference data set using CuBlock and ComBat while removing the groups *B* and *D* from the platforms HUG and AG1. In other words, these two platforms were normalized only with *A* and *C* samples. The biological difference between *A* and *C* is that 25% of *C* is made of *B* RNA samples. The other four platforms were normalized with all four biological groups. As it can be seen in Figure 7A,C, CuBlock results in the clustering of the *C* samples of HUG and AG1 separately and closer to the *D* cluster of the other platforms than to their *C* cluster. However, the *A* cluster remains a well-defined cluster for all platforms. This suggests that, in the two platforms with missing groups, CuBlock emphasizes the difference between the available data. For HUG and AG1, this means emphasizing the differences between *A* and *C* (the only groups it sees), thus bringing *C* closer to the *D* cluster formed by the other platforms, since, as *C* itself, *D* is also a mixture of *A* and *B*. When using ComBat (Figure 7B,D), the *C* samples of HUG and AG1 are even more mixed with the *D* samples, and the *A* samples of the two platforms tend to be closer (in the *t*-SNE plot) to the *C* samples of the four other platforms, with some of them even being clustered (in the dendrogram) in this group.

**Fig. 7.**
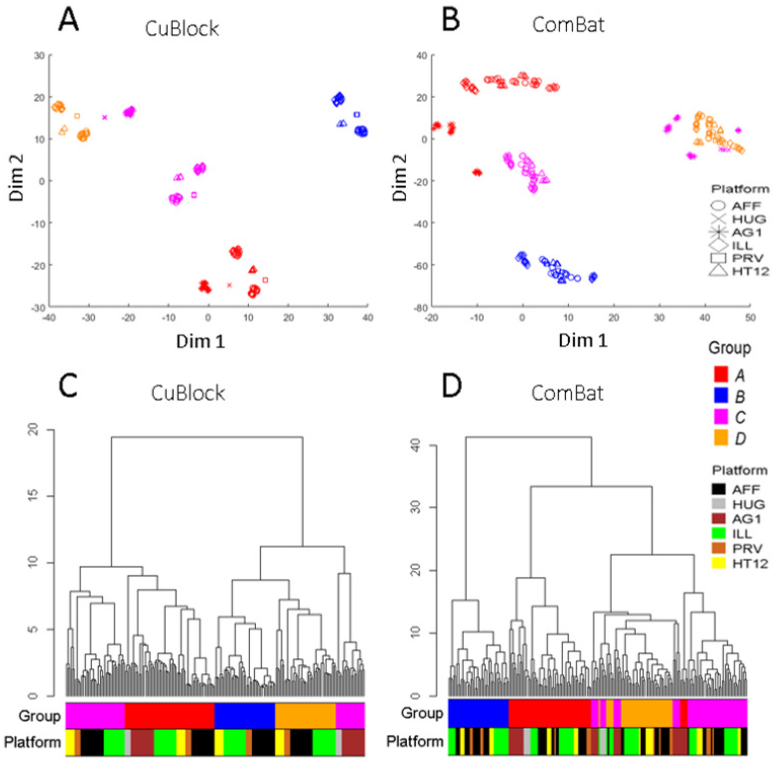
t-SNE dimension reduction and dendrogram plots for the reference data set, with exclusion of the *B* and *D* samples from platforms HUG and AG1, after normalization with CuBlock and ComBat. Point color and shape indicate biological group and platform, respectively (right-hand legend). A: t-SNE for CuBlock normalized data; perplexity (Prp) and mean silhouette index (SI) values (see Section 2.3.2): Prp = 15, SI = 0.80. B: corresponding analysis for ComBat-normalized data; Prp = 10, SI = 0.67. C, D: Dendrograms for CuBlock (C) and ComBat (D) normalized data; color bars under the dendrograms indicate the biological group and platform corresponding to each leaf.

## 5 Conclusion

We have introduced an algorithm for cross-platform normalization of geneexpression microarray data as well as a strategy to validate cross-platform normalization methods, with a focus on the capacity of the algorithm to properly separate samples from different biological groups after normalization and across multiple platforms. Overall, CuBlock showed good results on the two data sets used in this evaluation, a data set specifically generated for testing and standardization purposes and a data set from an actual experimental study. CuBlock could always differentiate, clearly, the underlying biological groups after mixing data from up to 6 different platforms. Nevertheless, we observed that within each biological group the algorithm tends to subcluster samples by platform, indicating a remaining, yet comparatively small, platform effect. Compared to simply taking the log_2_ transformation of the microarray data, CuBlock showed significant improvement in distinguishing the biological groups and mixing the platforms. The ComBat algorithm (Johnson *et al*., 2007) showed also good performance on the reference data set, with better mixing of platform data than all the other methods tested. However, on the experimental data set, where samples are from different individuals and the difference between biological groups might become less obvious than in the reference data set, ComBat did not perform as well. Platform mixing was still good but the distinction between the two biological groups was not clear. As mentioned in the introduction, ComBat also requires the platforms to be normalized together, making it a less convenient method for systematic application to multiple data sets. DBNorm was only used for the normalization of the experimental data set, as it proved computationally much more time demanding than the rest. Both DBNorm and YuGene performed only slightly betterthan log_2_. For the set evaluated, Shambhala lagged clearly behind the other methods, arguably including a simple log_2_ transformation.

The CuBlock transformation can be thought as remaining close to the input data. CuBlock fits cubic polynomials to data blocks that are found by *k*-means clustering, thereby trying to best fit the different distributions found in the data corresponding to a sample (different blocks need not have the same distribution) and it does so without assuming a shape for these distributions. As a consequence, CuBlock emphasizes the differences within the input data, making it sample-composition dependent despite the only step in the algorithm where the microarray samples from a given platform are considered together is when applying the *k*-means algorithm (afterwards, each sample is considered separately). This sample dependence was highlighted in this study when analyzing the reference data set after removing biological groups in some platforms. It is however less prominent than for algorithms normalizing platforms together, such as ComBat. In summary, we have shown that CuBlock can be applied to data from multiple microarrays in a platform agnostic way and preserves the biological grouping of the samples, demonstrating a good performance for different types of samples. It is therefore a tool appropriate for gene-expression studies based on multiple microarray sets collected along multiple platforms and at different times, thus facilitating the extraction of knowledge from the wealth of microarray data available in public repositories and enabling the use of these repositories as sources of Real-World Data.

## Acknowledgements

The authors wish to thank Malu Calle for useful discussions during the preparation of the manuscript.

