## Supplementary Information for "CuBlock: A cross-platform normalization method for gene-expression microarrays"

October, 2020

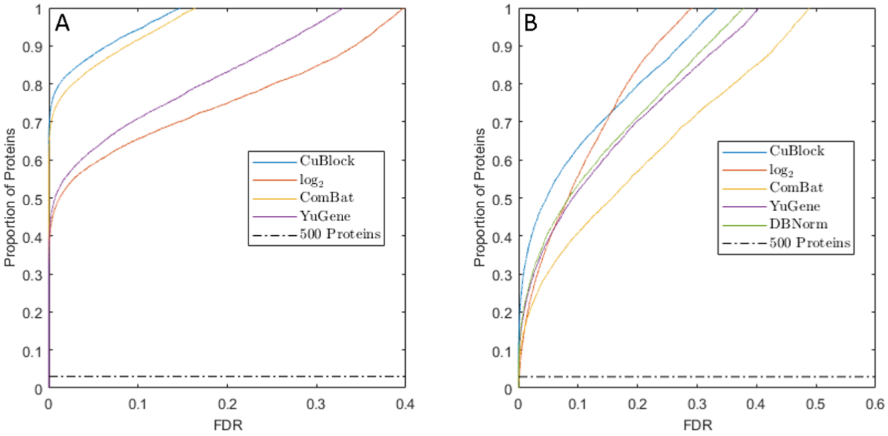


**Fig. S1:** ROC-like curves. The cumulative distribution function of the FDR-adjusted $p$-values (also called $q$-values) plotted for the reference and the experimental data set after normalization with CuBlock, $\log_2$, ComBat, YuGene and DBNorm. The dotted horizontal line represents the 500-protein cut. Plot A corresponds to the reference data set. Plot B corresponds to the experimental data set.

```
GetTargetValues(BS)
BS is a sorted array with mean 0 and standard deviation 1.

    L ← line of same size as BS between − 1 and 1 with equidistant points
    iStdUp ← the index corresponding to the point in BS which is the closest to 1
    iStdDown ← the index corresponding to the point in BS which is the closest to − 1
    FOR p ∈ [3,5,7,9,11,13,15,17,21]
      D ← L^p
      IF mean(D[iStdDown: iStdUp]) IS SMALLER than 0.1
        RETURN D
      END IF
    END FOR
    END FUNCTION
```

**Fig. S2:** Pseudocode describing the GetTargetValues algorithm called in the CuBlock algorithm (Figure 1 in the main text). An example illustration is given in Figure S5.

**ModPol**$(B, P)$
$B$ are the block values.
$P$ is the polynomial previously fitted to the block $B$.

$n \leftarrow$ size of $B$
$BS \leftarrow$ sort $B$ in ascending order
$\widehat{BS} \leftarrow$ evaluate the polynomial $P$ at $BS$
IF1 $\widehat{BS}$ ISNOT sorted in ascending order
  $[xDown1, iDown1] \leftarrow$ get the first value and corresponding index in $\widehat{BS}$ where the array decreases
  $[xDownL, iDownL] \leftarrow$ get the last value and corresponding index in $\widehat{BS}$ where the array decreases
  $[xUp1, iUp1] \leftarrow$ get the first value and corresponding index in $\widehat{BS}$ where the array increases
  $[xUpL, iUp] \leftarrow$ get the last value and corresponding index in $\widehat{BS}$ where the array increases
  IF2 $\widehat{BS}[iUp1]$ IS EQUAL to $\widehat{BS}[1]$ AND $\widehat{BS}[iUpL]$ IS EQUAL to $\widehat{BS}[n]$
    IF3 the number of points between the first index and $iDown1$ ISBIGGER than the number of point between $iDownL$ and the last index $n$
      $\widehat{BS}[iDown1: iDownL] \leftarrow xDown1$
      $\widehat{BS}[(iDownL + 1): n] \leftarrow xDown1 + \widehat{BS}[(iDownL + 1): n] - xDownL$
    ELSE3
      $\widehat{BS}[iDown1: iDownL] \leftarrow xDownL$
      $\widehat{BS}[1: (iDown1 - 1)] \leftarrow xDownL + \widehat{BS}[1: (iDown1 - 1)] - xDown1$
    END IF3
  ELSE2
    $\widehat{BS}[iUpL: n] \leftarrow xUpL$
    $\widehat{BS}[1: iUp1] \leftarrow xUp1$
  END IF2
END IF1
$\hat{B} \leftarrow$ sort $\widehat{BS}$ back in original order of $B$
END FUNCTION

**Fig. S3:** Pseudocode describing the ModPol algorithm called in the CuBlock algorithm (Figure 1 in the main text). An example illustration is given in Figure S6.

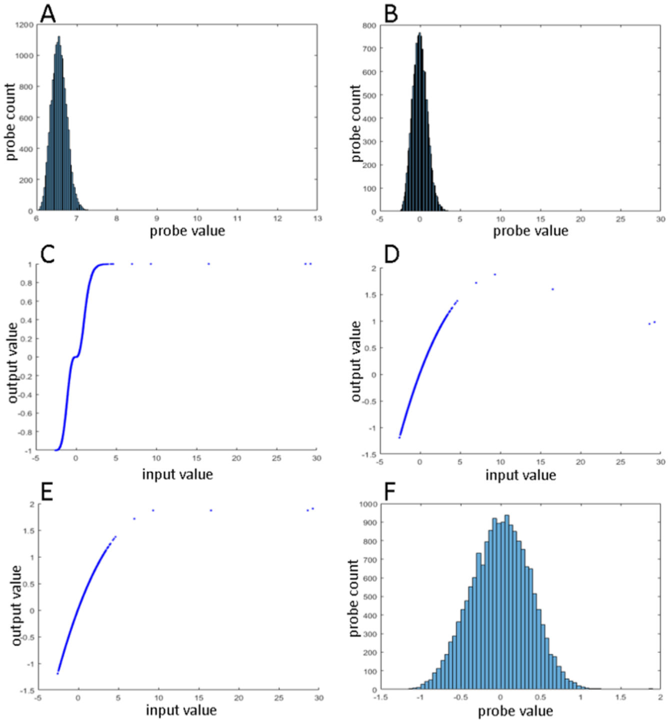


**Fig. S4:** The different steps of the CuBlock algorithm for an example block. A: histogram of the untransformed block. B: histogram of the z-transformed block. C: target values (output values) of the sorted z-transformed block (input values) as obtained after the GetTargetValues algorithm. D: values after evaluation of the fitted polynomial (output value) on the sorted z-transformed block (input value); the cubical polynomial's coefficients are such that these values minimize the mean root square error with the target values plotted in C. E: modified values (output value) as obtained after the ModPol algorithm; the decreasing values in D are corrected in order to preserve data sorting upon normalization. F: histogram of the normalized block.

The example block contains a few positive outliers that are identified by the fitted cubical polynomial in the decreasing part; this part is corrected by equating it to the last increasing point and letting the final increasing part continue its growth from this point.

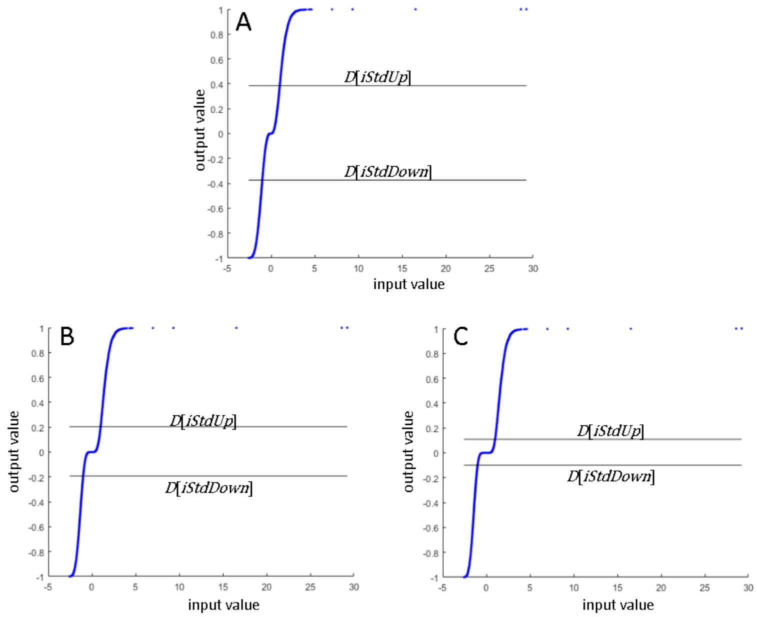


**Fig. S5:** Illustration of possible target values (output value) with respect to the sorted z-transformed block (input value) as described in the GetTargetValues algorithm. $D$ is a vector containing the target values and equals $L^p$ where $p$ is an uneven power and $L$ is a vector of equidistant points between $-1$ and 1 (the length of $L$ is the number of points in the block). $[iStdDown : iStdUp]$ represents the set of indices corresponding to the points in the sorted z-transformed block that are between $-1$ and 1, i.e. the points within standard deviation; mean$(D[iStdDown : iStdUp])$ is the average of the output values which are contained between the two horizontal lines in the plots. The target values are $D = L^p$ where $p$ is the smallest uneven power such that mean$(D[iStdDown : iStdUp])$ is smaller than 0.1.
A: target values for $p = 3$; mean$(D[iStdDown : iStdUp]) = 0.095$. B: target values for $p = 5$; mean$(D[iStdDown : iStdUp]) = 0.033$. D: target values for $p = 7$; mean$(D[iStdDown : iStdUp]) = 0.013$.
In this example block, the chosen $p$ is 3.

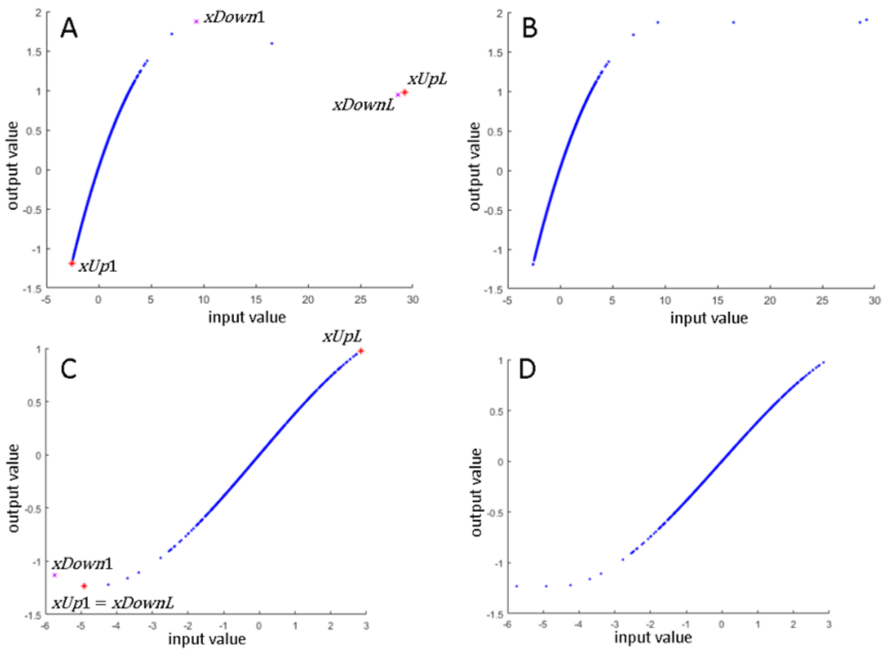


**Fig. S6:** Illustration of example modifications to correct for the decreasing values in the evaluation of the fitted polynomial (output value) on the z-transformed sorted block (input value) as described in the ModPol algorithm. A-B: example on one block with the output values before (A) the modifications and after (B). C-D: example on another block with the output values before (C) the modifications and after (D).
$xDown1$, resp. $xDownL$, is the first, resp. last, decreasing value and $xUp1$, resp. $xUpL$, is the first, resp. last, increasing value. These values are used to identify the decreasing part(s) which will be modified to be equal to the closest point in the main increasing part, i.e. the part containing most of the points.

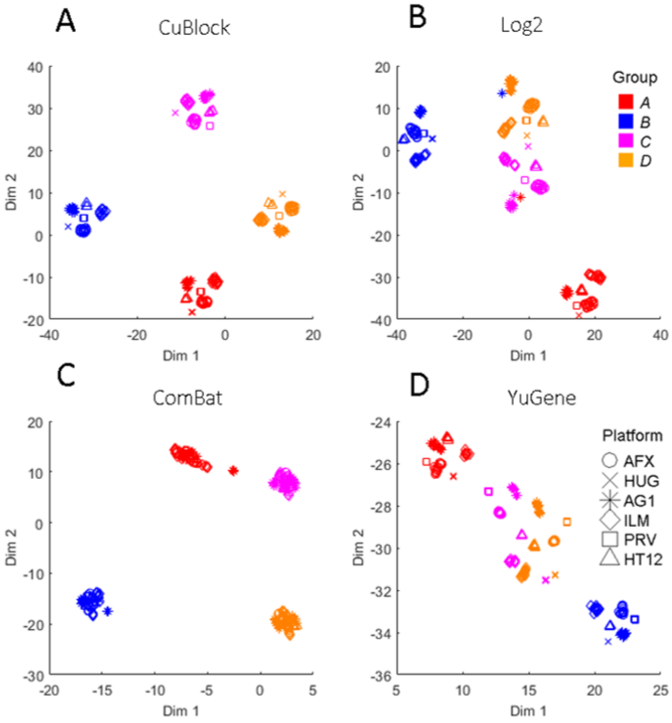


**Fig. S7:** $t$-SNE dimension reduction of the reference data set after normalization with CuBlock, $\log_2$, ComBat and YuGene. A: $t$-SNE for CuBlock normalized data; point color and shape indicate biological group and platform, respectively (right-hand legend); perplexity (Prp) and mean silhouette index (SI) values (see Section 2.3.2): Prp = 25, SI = 0.97. B: $t$-SNE for $\log_2$-normalized data; Prp = 20, SI = 0.74. C: $t$-SNE for ComBat-normalized data; Prp = 45, SI = 0.96. D: $t$-SNE for YuGene-normalized data; Prp = 78, SI = 0.68.

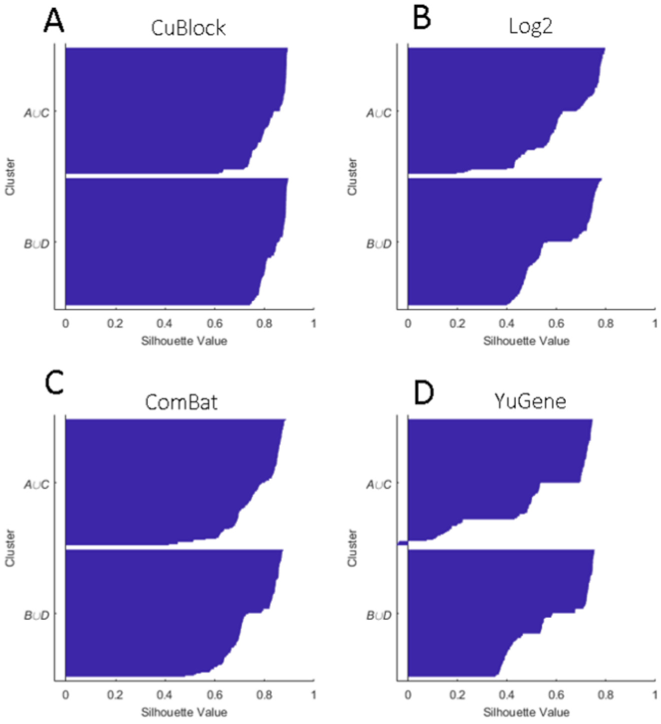


**Fig. S8:** Silhouette plots of the reference data set after normalization with CuBlock, $\log_2$, ComBat and YuGene. The given clusters are the groups $A \cup C$ and $B \cup D$. A: silhouette plot for CuBlock-normalized data; mean silhouette index (SI) values per group: 0.83 ($A \cup C$) and 0.84 ($B \cup D$). B: silhouette plot for $\log_2$-normalized data; SI values: 0.64 ($A \cup C$) and 0.61 ($B \cup D$). C: silhouette plot for ComBat-normalized data; SI values: 0.77 ($A \cup C$) and 0.75 ($B \cup D$). D: silhouette plot for YuGene-normalized data; SI values: 0.53 ($A \cup C$) and 0.59 ($B \cup D$).

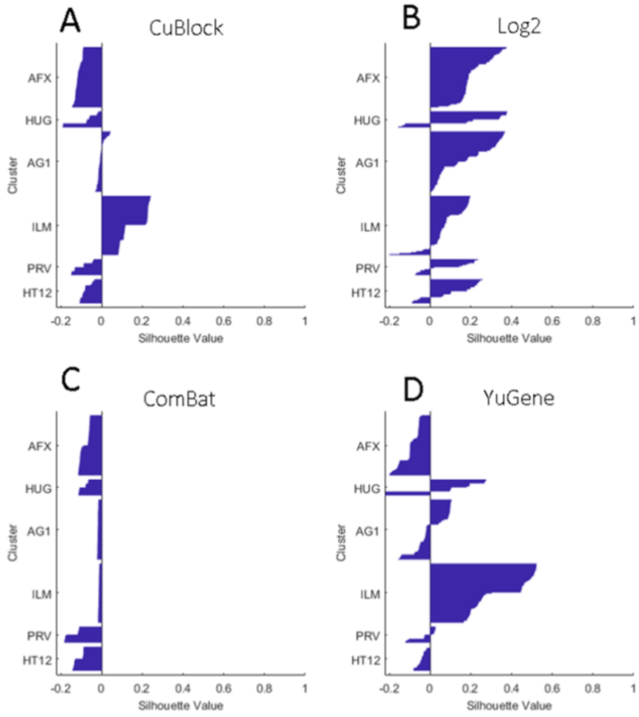


**Fig. S9:** Silhouette plots of the reference data set after normalization with CuBlock, $\log_2$, ComBat and YuGene. The given clusters are the platforms. A: silhouette plot for CuBlock-normalized data; mean silhouette index (SI) values per platform: -0.12 (AFX), -0.09 (HUG), 0.00 (AG1), 0.16 (ILM), -0.10 (PRV) and 0.08 (HT12). B: silhouette plot for $\log_2$-normalized data; SI values: 0.22 (AFX), 0.19 (HUG), 0.17 (AG1), 0.08 (ILM), 0.08 (PRV) and 0.11 (HT12). C: silhouette plot for ComBat-normalized data; SI values: -0.08 (AFX), -0.09 (HUG), -0.02 (AG1), -0.01 (ILM), -0.15 (PRV) and -0.11 (HT12). D: silhouette plot for YuGene-normalized data; SI values: -0.10 (AFX), 0.09 (HUG), 0.01 (AG1), 0.35 (ILM), -0.03 (PRV) and -0.04 (HT12).

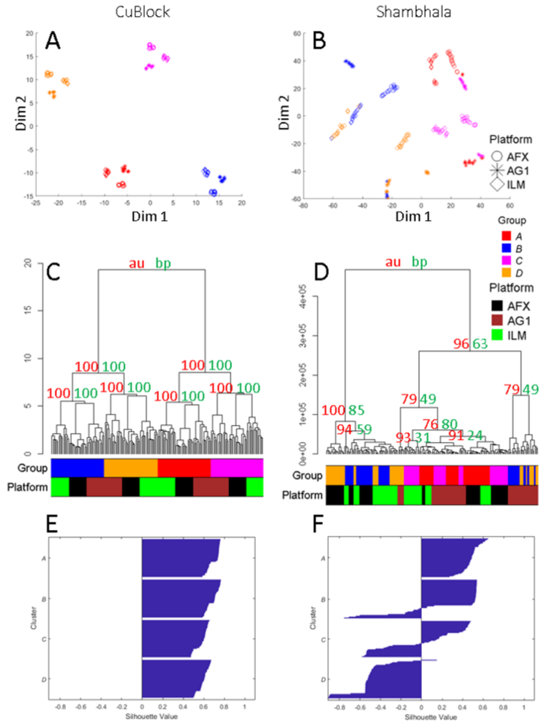


**Fig. S10:** *t*-SNE dimension reduction, dendrogram and silhouette plots for the reference data set (three platforms) after normalization with CuBlock and Shambhala. A: *t*-SNE for CuBlock normalized data; point color and shape indicate biological group and platform, respectively (right-hand legend); perplexity (Prp) and mean silhouette index (SI) values (see Section 2.3.1): Prp = 25, SI = 0.97. B: corresponding analysis for Shambhala-normalized data; Prp = 5, SI = 0.28. C, D: dendrograms for CuBlock (C) and Shambhala (D) normalized data; color bars below the dendrograms indicate the biological group and platform corresponding to each leaf; the BP (green) and AU (red) values (see Section 2.3.3) for some selected clusters are indicated at the origin of the branches. E: silhouette plot for CuBlock-normalized data, using the groups $A$, $B$, $C$ and $D$ as given clusters; SI values: 0.70 ($A$), 0.69 ($B$), 0.58 ($C$) and 0.60 ($D$). F: silhouette plot for Shambhala-normalized data; SI values: 0.49 ($A$), 0.27 ($B$), 0.03 ($C$) and -0.52 ($D$).

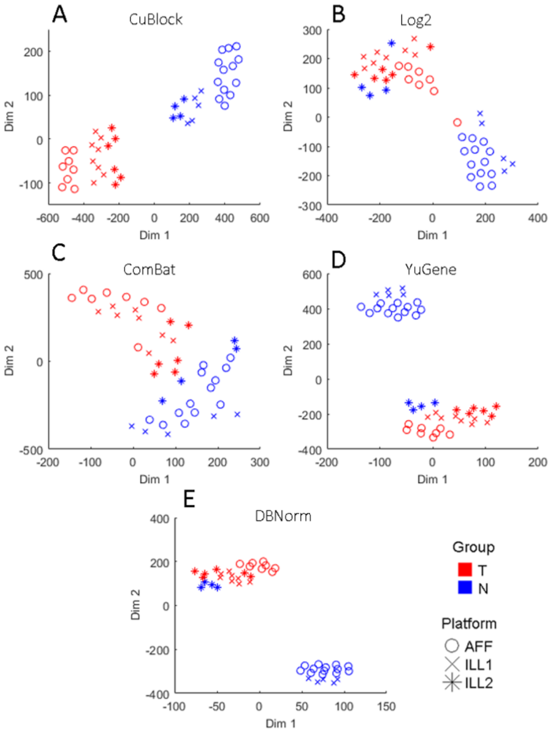


**Fig. S11:** $t$-SNE dimension reduction of the experimental data set after normalization with CuBlock, $\log_2$, ComBat, YuGene and DBNorm. A: $t$-SNE for CuBlock normalized data; point color and shape indicate biological group and platform, respectively (right-hand legend); perplexity (Prp) and mean silhouette index (SI) values (see Section 2.3.2): Prp = 10, SI = 0.93. B: $t$-SNE for $\log_2$-normalized data; Prp = 15, SI = 0.61. C: $t$-SNE for ComBat-normalized data; Prp = 10, SI = 0.62. D: $t$-SNE for YuGene-normalized data; Prp = 15, SI = 0.75. E: $t$-SNE for DBNorm-normalized data; Prp = 15, SI = 0.75.

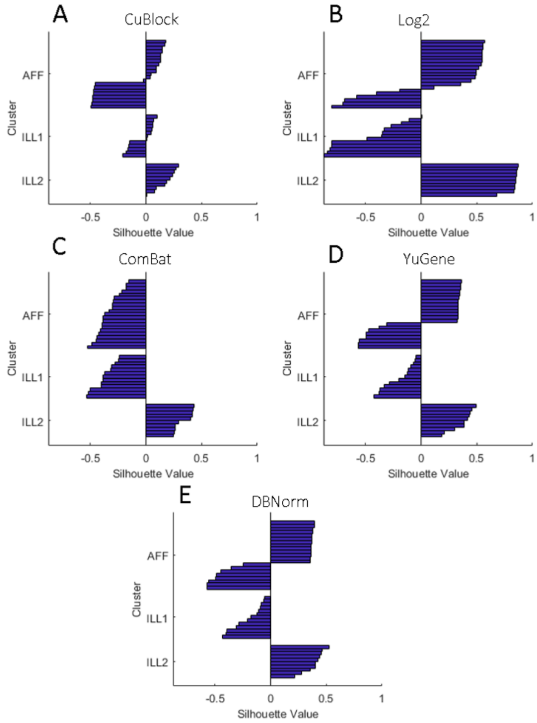


**Fig. S12:** Silhouette plots of the experimental data set after normalization with CuBlock, $\log_2$, ComBat, YuGene and DBNorm. The given clusters are the platforms. A: silhouette plot for CuBlock-normalized data; mean silhouette index (SI) values per platform: -0.12 (AFF), -0.03 (ILL1) and 0.18 (ILL2). B: silhouette plot for $\log_2$-normalized data; SI values: 0.19 (AFF), -0.48 (ILL1) and 0.83 (ILL2). C: silhouette plot for ComBat-normalized data; SI values: -0.34 (AFF), -0.38 (ILL1) and 0.34 (ILL2). D: silhouette plot for YuGene-normalized data; SI values: 0.03 (AFF), -0.20 (ILL1) and 0.37 (ILL2). E: silhouette plot for DBNorm-normalized data; SI values: 0.05 (AFF), -0.21 (ILL1) and 0.40 (ILL2).
